# Exploring the Biological Mechanisms of Severe COVID-19 in the Elderly: Insights from an Aged Mouse Model

**DOI:** 10.1101/2024.09.24.614691

**Authors:** Li Ma, Xian Lin, Meng Xu, Xianliang Ke, Di Liu, Quanjiao Chen

## Abstract

Coronavirus disease 2019 (COVID-19), caused by severe acute respiratory syndrome coronavirus 2 (SARS-CoV-2), has resulted in a global health crisis, particularly affecting the elderly, who are more susceptible to severe outcomes. However, definitive parameters or mechanisms underlying the severity of COVID-19 in elderly people remain confused. Thus, this study seeks to elucidate the mechanism behind the increased vulnerability of elderly individuals to severe COVID-19. We employed an aged mouse model with a mouse-adapted SARS-CoV-2 strain to mimic the severe symptoms observed in elderly patients with COVID-19. Comprehensive analyses of the whole lung were performed using transcriptome and proteome sequencing, comparing data from aged and young mice. For transcriptome analysis, bulk RNA sequencing was conducted using an Illumina sequencing platform. Proteomic analysis was performed using mass spectrometry following protein extraction, digestion, and peptide labeling. We analyzed the transcriptome and proteome profiles of young and aged mice and discovered that aged mice exhibited elevated baseline levels of inflammation and tissue damage repair. After SARS-CoV-2 infection, aged mice showed increased antiviral and inflammatory responses; however, these responses were weaker than those in young mice, with significant complement and coagulation cascade responses. In summary, our study demonstrates that the increased vulnerability of the elderly to severe COVID-19 can be attributed to an attenuated antiviral response and the overactivation of complement and coagulation cascades.

## 1. Introduction

Coronavirus disease 2019 (COVID-19), caused by the novel emerging severe acute respiratory syndrome coronavirus 2 (SARS-CoV-2), has triggered a worldwide health emergency, resulting in up to 6.9 million confirmed deaths (as of May 1, 2024). Age, sex, and comorbidities are significant risk factors associated with COVID-19 mortality and determine the most effective treatment strategies for high-risk patients (Laxminarayan et al., 2020). The risk of hospitalization and death rises with age, with the fatality rate increasing from 2.3% in the general population to 14.8% in patients over 80 years old (Wu and McGoogan, 2020). Studies revealed the highest susceptibility to infection among older individuals (Wu et al., 2020b, Zhang et al., 2020). Throughout the COVID-19 pandemic, elderly human populations exhibit higher morbidity and mortality rates (Vestergaard et al., 2020). Similarly, animal models show that SARS-CoV-2 causes more severe clinical phenotypes in older animals than in their younger counterparts (Kim et al., 2022). These findings demonstrated that age is the greatest risk factor for severe COVID-19.

Age is significantly correlated with COVID-19 severity; however, the underlying mechanisms remain unknown. Dysfunctional immune responses are crucial contributors to severe SARS-CoV-2 infections in the elderly population. Angiotensin-converting enzyme 2(ACE2) serves as a key receptor for SARS-CoV-2 infection (Liu et al., 2020). ACE2 expression may decrease with age, contributing to lung damage caused by SARS-CoV-2 in elderly people (Baker et al., 2021). Additionally, type I interferon response is impaired in elderly people, thus increasing their susceptibility to viral infection. The immune system undergoes various age-related changes in older adults, one of which is characterized by chronic, low-grade inflammation, known as inflammaging (Shive and Pandiyan, 2022). These preexisting low-grade inflammatory cytokines can create a more pro-inflammatory state in elderly patients. Another type of immune system dysfunction in the elderly is immune senescence. Marked changes can be observed in the immune system with aging, such as reduced migration, chemotaxis, and phagocytosis of neutrophils, macrophages, and dendritic cells. Others include diminished T-cell immune responses due to decreased expression of CD28 and the total numbers of CD4+ and CD8+cells (Crooke et al., 2019, Lee et al., 2022, Grubeck-Loebenstein et al., 2009). These changes in inflammaging and immune senescence appear to intensify during COVID-19, potentially increasing the disease severity.

Some correlations between aging and COVID-19 have been identified. However, definitive parameters or mechanisms underlying the severity of COVID-19 in elderly people remain unclear. The comprehensive profile of the host response to SARS-CoV-2 infection in young and aged mice at the transcriptome and proteome levels was still lacking. Therefore, this study aims to show the comprehensive profile of the host response and elucidate the mechanism behind the increased vulnerability of elderly individuals to severe COVID-19.

## 2. Materials and Methods

### 2.1. Virus and cells

An mouse-adapted SARS-CoV-2 (MAV) strain was obtained from Professor Meilin Jin of Huazhong Agricultural University (Huang et al., 2021). The MAV strain was cultured and quantified in Vero E6 cells using standard plaque-forming units techniques. Vero E6 cells were cultured in Dulbecco’s Modified Eagle Medium (DMEM) medium supplemented with 10% FBS and incubated at 37°C incubator with 5% CO2. All experiments involving viral infections were conducted in a Biosafety Level 3 laboratory at the Wuhan Institute of Virology.

### 2.2. Mouse experiment

The study followed the ARRIVE guidelines (https://www.nc3rs.org.uk/arrive-guidelines). Three-week-old and 18-month-old SPF C57BL/6j mice were purchased from Beijing Vital River Laboratory and Jackson Laboratory, respectively. All mice were cultured in a specific-pathogen-free culture environment. They were then randomly divided into four cohorts using a random number generator, each consisting of nine mice: young and aged groups without infection and young and aged groups with infection. Each mice was fed, monitored and sacrificed in the same way to minimise potential confounders. Anesthesia was induced in the mice through intraperitoneal administration of tribromoethanol at a dose of 250 mg/kg body weight. Mice in the infected groups were intranasally inoculated with 2 × 10^5^ plaque-forming units of MAV in 30 μL of DMEM per mouse, whereas mice in the uninfected groups were administered 30 μL of DMEM per mouse. Daily monitoring of weight changes continued for all mice until the end of the study. Based on transcriptome and proteome sequencing targets, the minimum biological replication was 3. Lung samples were collected from 25 mice. In details, 3 mice in each group at three days post-infection for proteomic analyses and 3 young, 3 aged, 4 MAV infected young, and 3 MAV infected aged mice at three days post-infection for transcriptomic analyses.

### 2.3. RNA extract

The lungs were homogenized in TRIzol reagent and then centrifuged at 12,000 rpm for 15 min at 4°C. The resulting supernatant was collected, and the total RNA was extracted using the phenol-chloroform method following the instructions of the manufacturer..

### 2.4. Immunohistochenmical analysis

The lungs were excised and fixed in 4% paraformaldehyde, subsequently embedded in paraffin and sectioned at 5 µm, and subsequently dried overnight at 42℃ before staining. Lung sections were deparaffinized, rehydrated using ethanol gradients, and underwent antigen retrieval with EDTA (pH 8.0) in a microwave oven. After blocking with 5% bovine serum albumin, tissue sections on slides were incubated with specific antibodies targeting the proteins of interest. For indirect immunofluorescence assay (IFA), fluorescent-labeled secondary antibodies were employed. Antibodies against C3, C4, and C5a were sourced from ProteinTech, and the slides were scanned using a Pannoramic MIDI system (3DHISTECH).

### 2.5. Western Blotting

The lungs were homogenized in protein extract buffer with protease inhibitor cocktail (MCE) at 4°C and then clarified via centrifugation at 12,000 rpm. Protein concentration was determined by BCA-based approach. After denaturation at 95°C for 5 min, equal amounts of protein (60 μg) from each group were separated using SDS-PAGE and transferred onto polyvinylidene difluoride membrane, by blocked with 5% non-fat milk at 25°C for 1 h. The membrane was washed with TBS buffer with 0.1% Tween-20. Then, the membranes were incubated with indicated antibodies at 4℃ overnight, followed with TBS-T washes. The membranes were then incubated with indicated secondary antibodies at room temperature. After three washes, the immunoblots were visualized using a ChemiDoc MP system (Bio-Rad). Protein ladder 26616 thermo was used as maker.

### 2.6. Next-Generation Sequencing Library Preparation and Sequencing

A comprehensive analysis of the lung tissue transcriptome was conducted. RNA quantification was performed individually using the Qubit 3.0 system. Samples meeting the quality standards underwent mRNA enrichment using a Dynabeads mRNA Purification Kit, followed by fragmentation, reverse transcription, and synthesis of double-stranded cDNA. Subsequently, adapters were attached to repair DNA ends, PCR products were cyclized, and libraries were prepared using the MGIEasy mRNA Library Prep Kit. RNA integrity was evaluated using a Bioanalyzer 2100 system. Libraries were sequenced using the Illumina NovaSeq.

### 2.7. Raw Data Processing

The raw data initially underwent quality control using fastp (Chen et al., 2018) (v0.20.1), where 10 low-quality bases were removed from the beginning and end of sequences. Sequences with <20 bases were discarded, and reads containing >6 N bases were removed. Subsequently, the resulting clean reads were mapped to the host and viral genomes by hisat2 (Kim et al., 2015) (v2.1.0). The mouse GRCm38 genome served as the host reference sequence, while the wild-type reference genome was utilized for SARS-CoV-2 (NC_045512.2). Samtools (Danecek et al., 2021) (v1.9) was employed for file format conversion and mapping rate statistics. A gene count matrix was generated using featureCounts (Liao et al., 2014) (v1.6.3) for subsequent analysis. Following this, fragments per kilobase of transcript per million mapped reads (FPKM) were computed, considering the gene length and mapped read count. FPKM quantifies gene expression levels by integrating sequencing depth and gene length in the calculation, making it a widely adopted method in current research.

### 2.8. RNA-seq data analysis

Differential expression analyses were performed using the DESeq2 (Love et al., 2014) package. Initially, raw counts were normalized based on the library size (using the rlog function) and transformed into log2 values. Subsequently, the sample batch effects were assessed via surrogate variable (SV) analysis using the SV package. Significant SVs were identified and incorporated into the model for subsequent differential analyses. Principal component analysis (PCA) was conducted using the prcomp function on the complete transcript datasets to visualize the clustering and variance of biological replicates across different conditions. Transcripts demonstrating fold changes > 1.5 and adjusted p-values corrected using the Benjamini–Hochberg (BH) ≤ 0.05 were considered statistically significant.

The log2-transformed counts were normalized using the remove BatchEffect function from the limma (Ritchie et al., 2015), incorporating identified SVs. Subsequently, the data were transformed into median-centered z-scores. These z-scores were then utilized for k-means clustering of all transcripts, with the number of clusters (k=10) determined based on the sum of the squared error and Akaike information criterion. Transcripts with similar expression patterns were grouped into clusters. The resulting clusters and differential expression of transcripts were visualized using heat maps created by ComplexHeatma package (Gu et al., 2016).

### 2.9. Protein preparation

To prepare samples, lung tissues from mice were homogenized, and the protein concentration was determined using a BCA assay kit. Equal amounts of protein from each sample were digested with lysis buffer, and TCA was added to precipitate the proteins. The samples were centrifuged at 4500 ×*g* for 5 min. Following that, the supernatant was discarded, and the pellet was washed 2–3 times with pre-cooled acetone. The dried pellet was then resuspended in 200 mM TEAB and sonicated to disperse the precipitate. Trypsin was added at a 1:50 enzyme-to-protein ratio (w/w) for overnight digestion. Subsequently, Dithiothreitol was included at a final concentration of 5 mM for reduction at 56°C for 30 min. Next, iodoacetamide was added at a final concentration of 11 mM and the mixture was incubated for 15 min at room temperature in the dark.

Peptides obtained from trypsin digestion were desalted using Strata X C18 columns (Phenomenex) and lyophilized via vacuum centrifugation for TMT labeling. The peptides were then reconstituted in 0.5 M TEAB and labeled following the guidelines of the manufacturers of the TMT reagent kit. Peptide fractionation was conducted using high-pH reverse-phase HPLC with an Agilent 300Extend C18 column, and liquid chromatography separation was performed using the UltiMate 3000 RSLCnano system.

### 2.10. Database search and proteomic analysis by LC-MS/MS

Raw MS files were analyzed using MaxQuant version 1.6.0.16 against the Uniprot mouse database, with a mass tolerance of 10 ppm for MS1 and 0.02 Da for MS2 searches. A 1% false discovery rate was applied for proteins and peptides, and only peptides containing more than six amino acid residues were considered for identification and quantitation. Label-free quantitation was conducted using the MaxLFQ algorithm within the MaxQuant software. The significance of the changes in protein abundance was assessed using a two-sided Student’s t-test with BH correction for the intensity of protein LFQ. Proteins identified by at least one unique peptide and two or more total peptides, with a p-value < 0.01, were considered statistically significant.

### 2.11. Enrichment of Gene Ontology (GO) and Kyoto Encyclopedia of Genes and Genomes database (KEGG) analysis

GO and KEGG analyses were performed using the whole lung transcriptome or all identified proteins as background, employing a two-tailed Fisher’s exact test. Correction for various hypothesis testing of p-values was conducted using clusterProfiler (Yu et al., 2012) (v3.14.3), applying the Benjamini–Hochberg false discovery rate method. Only significantly altered biological process GO terms, or KEGG pathways with BH corrected p< 0.05 were retained. The z-score concept, as described by the GO plot (Walter et al., 2015) (v1.0.2), was used to determine whether biological processes were more likely to decrease (negative value) or increase (positive value).

### 2.12. GO and KEGG interaction pathway network

The network was constructed by linking pathways with shared differentially expressed genes (DEG) (Jaccard similarity > 0.2) and clustering pathways with similar functional annotations. Nodes in the network were color-coded based on their representation in the k-means clustering analysis, and node size reflected the number of genes annotated in each pathway. Pathway networks were visualized in Cytoscape (Shannon et al., 2003) using a modified configuration from Metascape (Zhou et al., 2019).

### 2.13. Enrichment-based Clustering

To further cluster proteins based on their functions, enriched categories with p values < 0.05 were collected and transformed using -log10. These transformed values were then z-scored for each category. The z-scores were clustered using hierarchical clustering in Genesis and visualized as a heatmap in R using the gplots package (v3.1.3.1).

### 2.14. Statistical Analysis

Statistical analyses were performed in the R environment employing t-tests, Dunn’s Kruskal– Wallis tests, and Dunnett’s tests to determine statistical significance. A significance level of BH corrected p < 0.05 was considered statistically significant unless otherwise specified.

## 3. Results

### 3.1. Transcriptome and Proteome Analyses of Young and Aged Mice Lungs Infected with SARS-CoV-2

To elucidate the severe mechanism of COVID-19 in the elderly, we established an aged mouse infection model and compared whole-lung biological responses to SARS-CoV-2 between young and aged mice using transcriptome and proteome analyses. The animal experiments comprised four groups: young (3-week-old, 3W) mock infection (Y), young mice infected with MAV (YV), aged (19-month-old, 19M) mock infection (O), and aged mice infected with MAV (OV) groups. The animal infection experiment successfully reproduced the clinical symptoms observed in humans. Particularly, MAV infection caused severe disease phenotypes in aged mice but not in young mice (Detailed data on mouse infection are presented in a separate article that is being submitted, not shown in this article).

Overall, we used lung tissues from 13 mouse to do RNA sequencing: Y (n=3), O (n=3), YV (n=4), and OV (n=3). 32,770 RNAs were identified across the four groups. Genes with minimal deletions (<1 deletion per group) were selected for subsequent analysis, revealing 14,139 co-expressed RNAs, including 12,715 mRNA encoding proteins in the transcriptome sequencing (Fig. 1A). To produce proteomic data, we used lung tissues from 12 mouse: Y (n=3), O (n=3), YV (n=3), and OV (n=3). In total, 54,094 peptides were identified, containing 4,840 proteins. An overlap of 3,136 mRNA/genes and proteins was observed between the two omics datasets (Fig. 1B). Next, DEG and proteins (DEP) related to age and infection in the transcriptome and proteome data were analyzed. PCA of mRNA transcription and protein expression revealed clear clustering of samples within groups and distinct separation among groups (Fig. 1C). The overlapping genes/proteins reflected sample diversity (Supplementary Fig. S1A), with samples distinctly clustered based on RNA and protein profiles (Fig. 1C), indicating high-quality sample preparation. GO enrichment analysis revealed the central roles of these overlapping genes/proteins in cellular physiological activities and their structural functions (Supplementary Fig. S1B). The DEG and DEP were visualized in heatmaps, showing consistent expression patterns with PCA results (Supplementary Fig. S1C and D).

**Fig. 1.**
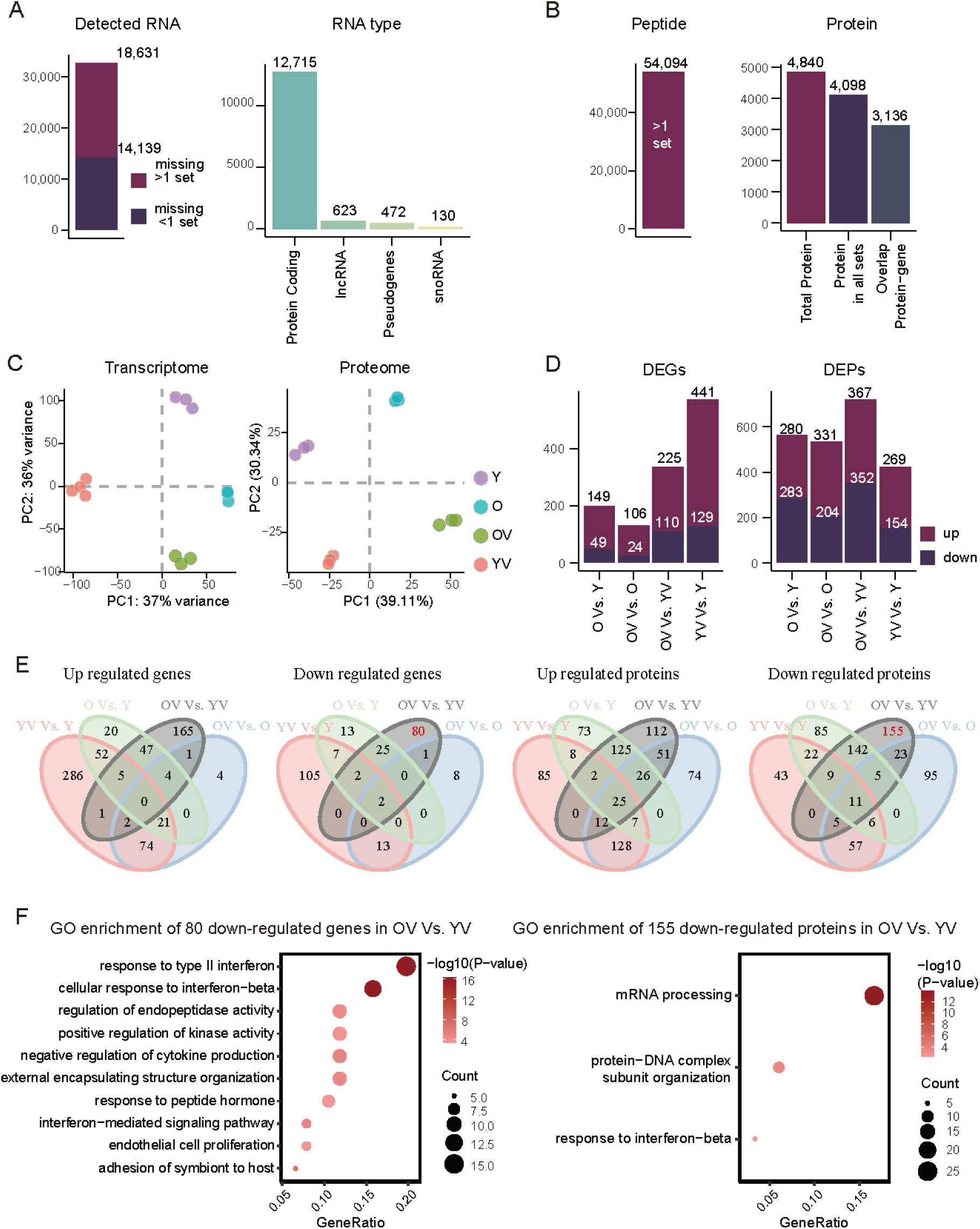
Comprehensive analysis of gene and protein expression profiles in mouse lungs across different groups. (A) The number and the highest concentration of four types of detected RNA: “Missing <1 set” indicates that only one sample per group of mice failed to detect RNA, and “missing >1 set” indicates more than one sample per group failed to detect RNA. (B) The number of identified and quantified peptides and proteins. “>1 set” indicates peptides detected in more than one set. (C) Principal component analysis (PCA) plots of transcriptome and proteome datasets. (D) Differentially expressed genes (DEG) and proteins (DEP) between groups. (E) Venn diagrams showing upregulated and downregulated genes and proteins across different groups. Supplementary excel 1 shows the genes involved. (F) Gene ontology (GO) enrichment analysis of uniquely downregulated genes or proteins in the OV Vs. YV group. The y-axis represents the biological processes, x-axis represents the gene ratio, bubble size represents the count of genes, and color intensity indicates -Log10 (adjusted p-value).

Next, DEG analysis was conducted between the groups. Overall, 149 genes and 280 proteins were upregulated, while 49 genes and 283 proteins were downregulated compared to those in the Y group. Overall, 225 mRNA and 367 proteins were upregulated, while 110 mRNA and 352 proteins were downregulated compared to those in the YV group, indicating a further increase in the DEG and DEP upon virus infection (Fig. 1D).

To investigate the correlation between aging and severe COVID-19, Venn diagrams were used to identify the unique and commonly regulated DEGs. The OV group exhibited 165 specifically upregulated and 80 downregulated genes, along with 112 and 155 specifically upregulated and downregulated proteins, respectively, compared to those in the YV group (Fig. 1E). Functional enrichment analysis was performed to examine the functions of these unique and common DEGs (Fig. 1F and Supplementary Table S1). The result showed that in the OV group, 80 uniquely downregulated genes were enriched compared to those in the YV group in response to type II interferon and regulation of endopeptidase or kinase activity. Similarly, the 155 uniquely downregulated proteins were enriched in pathways related to mRNA processing, protein-DNA complex subunit organization, and response to interferon-beta, consistent with that of genes (Fig. 1F).

### 3.2. Attenuated Antiviral Response between Aged and Young Mice upon Virus Infection

The transcriptome analysis unveiled unique clusters of highly differentially expressed genes that were specific to each of the four mouse cohorts, highlighting distinct genetic profiles among the groups. Next, we explored the biological implications and significance of these genes. Using the top 1000 genes highly contributing to principal components 1 and 2, we performed k-means clustering, and two clusters were obtained. GO enrichment analysis was then performed for each cluster. For principal component 1, cluster 1 genes were predominantly expressed in the Y group and enriched in biological pathways related to cellular behavior such as cell migration, fate determination, and differentiation (Supplementary Fig. S2A). In contrast, cluster 2 genes were highly expressed in the OV group and enriched in immune-related pathways, including leukocyte migration, adhesion, and activation, and positive regulation of immune effector processes (Supplementary Fig. S2A). In principal component 2, cluster 1 genes were highly expressed in the YV group and involved in intracellular metabolism, activation of antiviral responses, and autophagy as mechanisms against injury and infection (Supplementary Fig. S2B). Cluster 2 genes, highly expressed in the O group, were significantly prominent in energy production, oxidation of organic compounds, and cellular respiration (Supplementary Fig. S2B). Distinct gene expression patterns were observed across the four groups.

To further elucidate the global patterns of gene expression and identify genes regulated by aging and SARS-CoV-2 infection, we conducted k-means clustering analysis on all DEGs in the following comparisons: O Vs. Y, OV Vs. O, YV Vs. Y, and OV Vs. YV. Ten clusters were identified, with cluster 9 positively correlated with viral infection, while clusters 4 and 8 showed negative correlations with infection. Aging positively correlated with clusters 1 and 5 but negatively correlated with cluster 6 (Fig. 2A). This correlation between aging and viral infection with these clusters (clusters 1, 4, 5, 6, 8, and 9) was further validated using an alternative method (gene expression) (Fig. 2B and Supplementary Fig. S2C). Functional enrichment analysis was then performed in 10 clusters, and scaled gene expression was visualized using a heatmap. The DEGs involved in interferon-alpha and interferon-gamma responses, as well as antigen presentation processes, are presented in a heatmap (Fig. 2C). The enriched GO terms within each cluster, along with the associated DEGs, were comprehensively depicted in the GO network (Supplementary Fig. S3). Specifically, GO enrichment analyses of clusters 1, 5, and 9—positively correlated with aging and viral infection, respectively,—revealed that cluster 9 was enriched in functions related to immune and inflammatory responses, defense against viruses, response to interferon-beta, and antigen presentation, confirming its association with viral infection (Fig. 2C and Supplementary Fig. S3). The data indicated that while the OV and YV groups activated antiviral immune pathways, aged mice exhibited attenuated innate antiviral responses to viral infection compared to young mice (Fig. 2B–D). Cluster 1 genes were enriched in functions related to cytoplasmic translation, antigen presentation, and immune activation, showing heightened expression in aged mice, particularly in the aged mock-infected mice (Fig. 2C). Cluster 5 genes were enriched in biological processes specifically upregulated in the OV group, including protein synthesis, structural organization, differentiation, metabolism, and axoneme assembly (Fig. 2C).

**Fig. 2.**
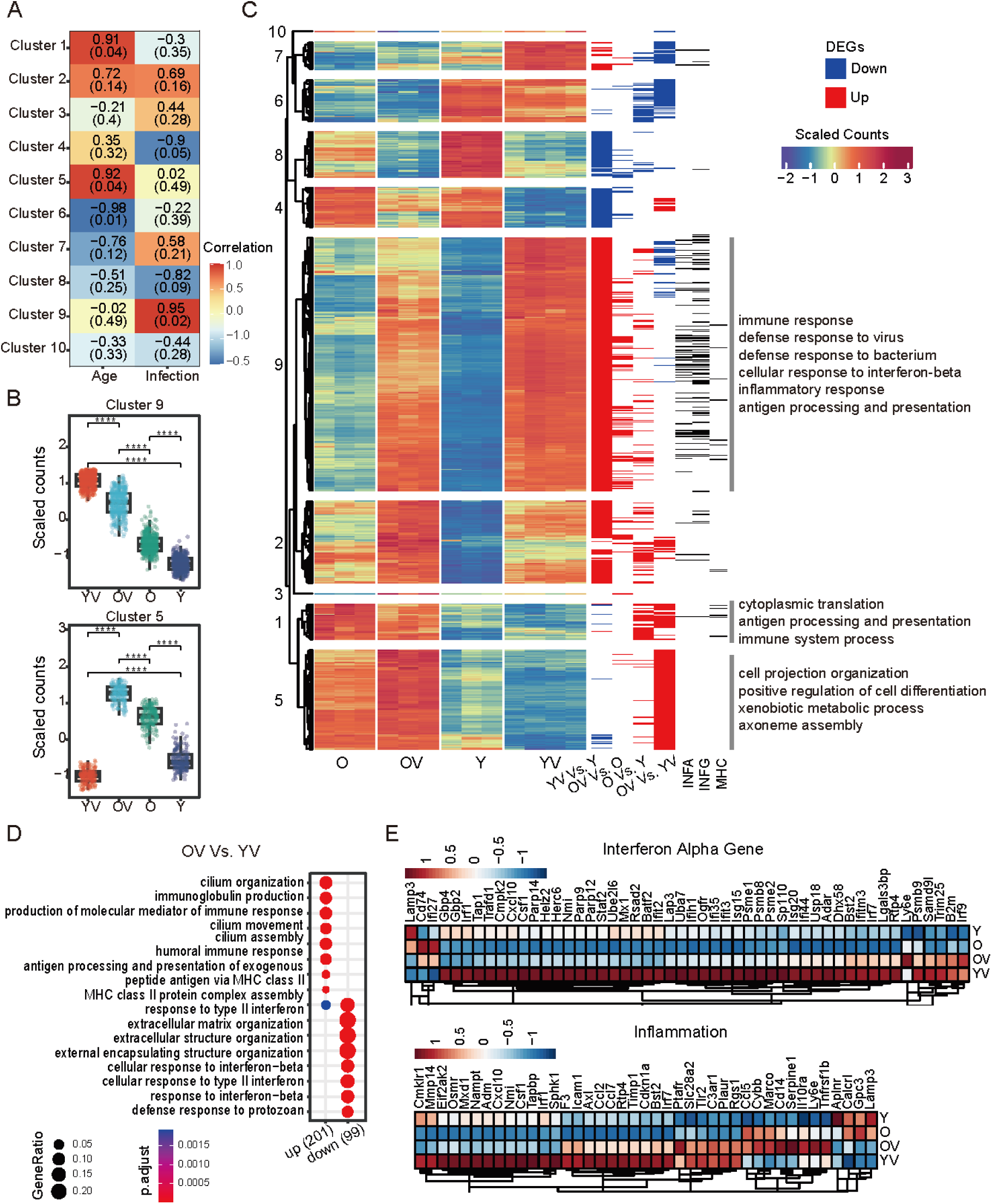
Gene expression profiling and functional annotation related to Severe Acute Respiratory Syndrome Coronavirus 2 (SARS-CoV-2) infection and age. (A) Heatmap showing a correlation between gene expression clusters and age/infection status. Cells represent correlation coefficients (red=positive, blue=negative) with corresponding p-values and statistical significance. (B) Box plots displaying gene expression in clusters 5 and 9, with asterisks denoting statistical significance (****p<0.0001). (C) Heatmap illustrating hierarchical clustering of DEGs across groups, with color intensity indicating expression levels. Sidebars denote up/downregulated genes in four groups based on Log2 Fold Change ≥1.5 or ≤-1.5 and p < 0.05. Black sidebars indicate gene pathways, with a prominent enriched pathway highlighted on the far right. (D) Bubble plot showing the functional enrichment analysis of differentially expressed genes (DEGs) between OV and YV groups. (E) Heatmap displaying highly variable genes (HVGs) in interferon alpha gene and inflammation pathways among the four groups, with color intensity indicating expression levels.

To further investigate the signaling pathways underlying poor prognosis in aged mice after viral infection, GO enrichment analysis was performed on DEGs, including OV Vs. O, YV Vs. Y, and OV Vs. YV. The results showed that the antiviral pathway was activated in aged and young mice after infection. However, the intensity was significantly weaker in aged mice than in young mice (Fig. 2D and Supplementary Fig. S4A). Next, to identify genes contributing to interferon and inflammatory response pathways, we analyzed highly variable genes (HVGs) among the groups. Consistent with the results of cluster 9, most of the HVGs were highly expressed in the YV group, including interferon-alpha genes such as Gbp4, Gbp2, and Irf1, as well as inflammation-related genes such as Eif2ak2, Osmr, and Mxd1. Differential gene activation patterns were observed in the OV group (Fig. 2E and Supplementary Fig. S4B). In summary, aged mice exhibit an attenuated antiviral immune response, potentially contributing to extensive tissue injury and a worse prognosis.

### 3.3. Overactivation of Complement and Coagulation Cascades in Aged Mice upon Virus Infection

In addition to the transcriptome analysis, mouse lung tissue underwent proteomic sequencing and analysis. Protein expression correlation analysis revealed high intragroup consistency and marked differences in protein expression profiles among the groups. The proteomic landscape of the O group positively correlated with that of the Y group, while a negative correlation was observed between the OV and YV groups (Supplementary Fig. S5A). To further explore the biological significance of these different protein expression profiles, differentially expressed proteomics (DEPs) were analyzed, and GO and KEGG enrichment analyses were conducted for O Vs. Y, OV Vs. O, YV Vs. Y, and OV Vs. YV. Enriched biological processes (p < 0.05) were collated. They were then clustered using one-way hierarchical clustering. In young mice, activation of antiviral immune responses post-infection, including signal transduction, interferon response, and inflammatory pathways, was observed and confirmed via transcriptome analysis. Furthermore, we found that the YV group exhibited a prominent acute inflammatory response, whereas the regulation of the inflammatory response was enriched in the O and OV groups. The OV group showed weaker antiviral immune responses after infection; however, a stronger humoral immune response, complement activation, and damage repair progress were observed in the YV group. These include wound healing, homeostasis, blood coagulation, protein activation cascade, and maturation (Fig. 3A). Further investigation from KEGG analysis revealed that the complement and coagulation cascades pathway was significantly enriched in the O (O Vs. Y) and OV (OV Vs. YV) groups (Fig. 3B and Supplementary Fig. S5B). We then identified 152 coregulated proteins from the intersecting DEGs of O Vs. Y and OV Vs. YV groups and performed KEGG enrichment analysis on these proteins (Fig. 3C). The complement and coagulation cascades pathway was remained enriched, comprising the greatest number of coregulated proteins (Fig. 3C). Furthermore, protein interaction network analysis was performed and two key biological pathways: “Complement and coagulation cascades” and “Regulation of Complement cascades” were identified (Fig. 3D). We then investigated the expression of complement-and coagulation-related proteins in mice, which were significantly prominent in the OV group (Fig. 3E). To further explore the expression of complement-and coagulation-related genes at the transcriptome level, the scaled gene expression is shown in a heatmap. Complement-and coagulation-related genes were highly differentially expressed in OV and YV groups (Fig. 3F).

**Fig. 3.**
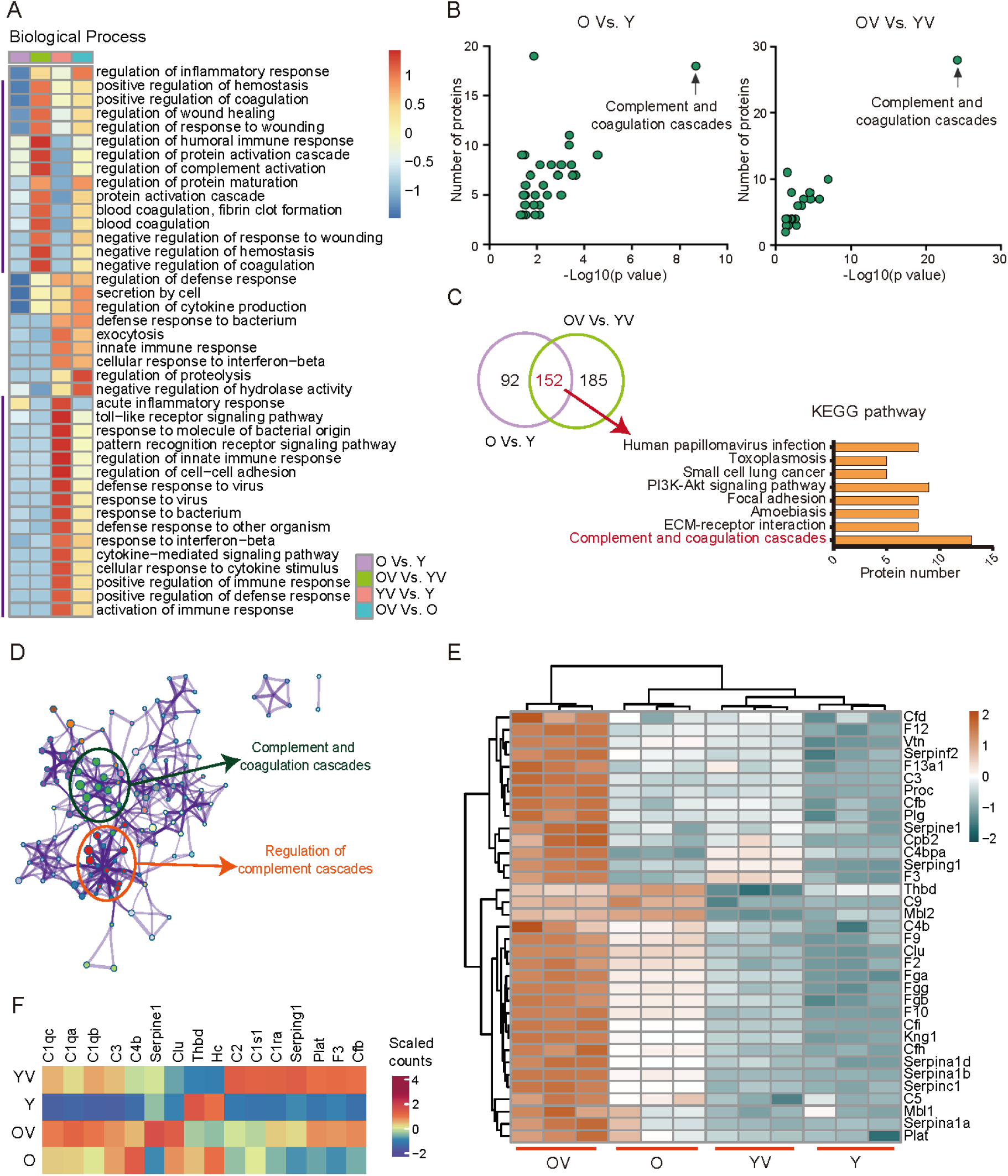
Protein expression profiling and functional annotation related to SARS-CoV-2 infection and age. (A) Heatmap depicting differences in activation of biological processes among groups: O Vs. Y, OV Vs. O, YV Vs. Y, and OV Vs. YV. Enrichment-based clustering details are provided in the Methods section. Red and blue colors indicate the extent of enrichment. (B) Scatter plots illustrating changes in protein quantity between groups O Vs. Y and OV Vs. YV. (C) Venn diagram displaying the intersection of shared datasets between comparisons: O Vs. Y and OV Vs. YV. Histogram showing enriched Kyoto Encyclopedia of Genes and Genomes database (KEGG) pathways of these DEPs. (D) Interaction network of proteins and KEGG pathways. Nodes represent proteins; edges indicate interactions and colors represent KEGG pathways. (E) Heatmap displaying expression of proteins in complement and coagulation cascades across groups: O Vs. Y, OV Vs. O, YV Vs. Y, and OV Vs. YV. The color shows scaled protein expression from blue (low) to orange (high). (F) Heatmap showing mean expression levels of genes in complement and coagulation cascades among groups: O Vs. Y, OV Vs. O, YV Vs. Y, and OV Vs. YV.

### 3.4. Complement and coagulation-associated proteins were increased in aged mice upon virus infection

As previously mentioned, the complement and coagulation pathway is activated in aged mice. To further verify the expression and activation of complement in the lung tissues of young and aged mice, IFA was performed. The complement system comprises approximately 40 proteins that coordinate the inflammatory response and possess direct antimicrobial properties. Activation occurs through three pathways (classical, lectin, and alternative), resulting in proteolytic processing of crucial components such as complement components 3, 4, and 5 (C3, C4, and C5) (Mathern and Heeger, 2015, Ricklin et al., 2010). Therefore, we assessed C3, C4, and C5, along with MBL-associated serine protease 2 (MASP2) in the lectin pathway. Following viral infection, C3, C4, C5, and MASP2 were found to be upregulated in young and aged mice. Particularly in the OV group, these components were significantly upregulated after infection. In the tissues of aged mice, we observed constitutively higher levels of C3, C4, C5, and MASP2 than in young mice, with an even greater increase in young infected mice (Fig. 4A). The results of western blotting analysis revealed higher levels of C3, C4, C5, and MASP2 in the O group than those in the Y group. The OV group showed the highest expression levels of C3, C4, C5, and MASP2, consistent with the IFA results (Fig. 4B).

**Fig. 4.**
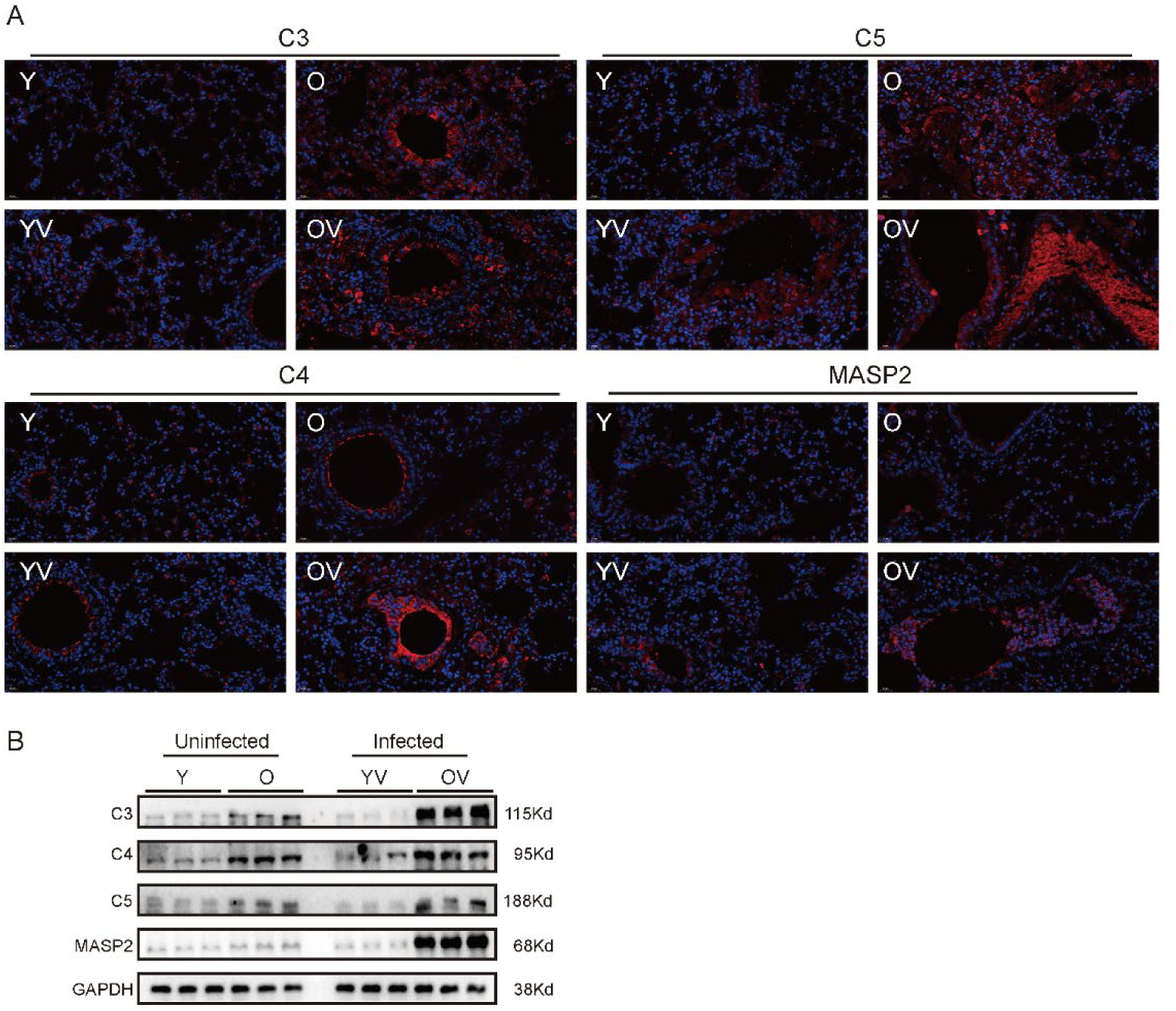
Expression of complement proteins in four groups. (A) Immunofluorescence assay depicting the location of complement proteins C3, C4, C5, and MBL-associated serine protease 2 (MASP2) in lung tissue from four groups: Y, O, YV, and OV. Complement proteins are shown in red, and cell nuclei in blue. (B) Western blot assay showing the expression levels of complement proteins C3, C4, C5, and MASP2 in lung tissues from four groups: Y, O, YV, and OV. GAPDH served as a loading control.

## 4. Discussion

This study aimed to investigate the mechanisms underlying severe COVID-19 in elderly individuals. Transcriptomic and proteomic analyses were conducted to identify DEGs and DEPs associated with aging and SARS-CoV-2 infection. The immune system of aged mice exhibited higher activation than that of young mice. Specifically, the expression of genes involved in antigen processing and presentation, xenobiotic metabolism, cell differentiation, and cytoplasmic translation were upregulated in aged mice. These findings were confirmed using transcriptomic and proteomic analyses. Upon SARS-CoV-2 infection, aged mice exhibited an activated antiviral immune response weaker than that in young mice, including responses related to inflammation, interferon-beta, and microbial defense. Processes protecting the body against infections and facilitating wound healing were boosted in aged mice and intensified after SARS-CoV-2 infection, particularly through the complement and coagulation cascade pathways. Following viral infection, we observed that aged mice exhibited a chronic inflammatory response involving the regulation of inflammatory response pathways, contrasting with the acute inflammatory response that is predominant in young mice.

Aging is characterized by chronic low-grade inflammation and immune system deterioration, leading to immunosenescence and reduced pathogen immunity, which heighten susceptibility to severe COVID-19 outcomes (Franceschi et al., 2018, Campisi et al., 2019, Bartleson et al., 2021). Studies consistently identify age as the primary risk factor for COVID-19 progression to acute respiratory distress syndrome and end-organ failure (Wu et al., 2020a, Bajgain et al., 2021, Ho et al., 2020). In aged mice, antibody-mediated depletion of self-renewing hematopoietic stem cells rejuvenates the immune system, enhancing adaptive immune responses to viral infections (Ross et al., 2024). Our study found that aged mice exhibited a diminished innate antiviral response post-infection, characterized by the reduced expression of genes involved in interferon signaling and antigen presentation, aligning with age-related immune decline.

The complement cascade enhances the ability of antibodies and phagocytic cells to clear microbes and damaged cells, while the coagulation cascade is crucial for blood clot formation and tissue repair (Ricklin et al., 2010, Furie and Furie, 2008). The complement system plays a dual role in host defense and inflammation; while crucial for pathogen elimination, uncontrolled complement activation can lead to tissue damage, thrombosis, and exacerbation of inflammation, which are hallmarks of severe COVID-19 (Levi et al., 2020, Tang et al., 2020b). Severe COVID-19 frequently manifests with coagulopathy, and disseminated intravascular coagulation is prevalent in most fatalities (Tang et al., 2020b, Huang et al., 2020, Chen et al., 2020). Anticoagulant therapy is associated with reduced mortality in patients with severe COVID-19 and sepsis-induced coagulopathy (Tang et al., 2020a). Patients with COVID-19 exhibit distinct complement activation than patients with influenza or non-COVID-19 respiratory failure (Ma et al., 2021). Systemic complement activation is also observed in children with multisystem inflammatory syndromes (Syrimi et al., 2021). The SARS-CoV-2 nucleocapsid proteins exacerbate lung injury via MASP2-mediated complement hyperactivation (Ali et al., 2021). Positive correlations between crucial COVID-19 in-hospital deaths and levels of C5a, C3a, and factor P, as well as serum IL-1β, IL-6, and IL-8, are observed (Alosaimi et al., 2021, Rockx et al., 2009). The inhibition of complement activation during coronavirus infection correlates with reduced systemic inflammation and tissue damage (Jiang et al., 2018, Gralinski et al., 2018, Yan et al., 2021, Rockx et al., 2009). Upregulated expression of complement and coagulation factors in the OV group may contribute to the pathogenesis of severe COVID-19 in the elderly population. Targeting complement signaling emerges as a promising treatment strategy for SARS-CoV-2 infection.

These findings suggest that aging involves intrinsic upregulation of complement components, which significantly intensifies upon viral infection, potentially exacerbating the severity of infections observed in elderly individuals. This heightened activation in elderly individuals may indicate a dysregulated immune response, underscoring the necessity for age-specific therapeutic approaches in managing viral infections.

Our findings enhance our understanding of how aging influences the immune response and viral pathogenesis. However, further research is needed to translate these findings into clinical interventions to reduce severe COVID-19 among elderly individuals. Developing age-specific therapeutic strategies, including increased antiviral therapy and modulation of the complement system, holds promise for improving clinical outcomes in this vulnerable population during the ongoing pandemic.

In conclusion, this study offers insights into age-dependent dysregulation of genes and cellular processes contributing to the severity of COVID-19. The antiviral immune response was activated after SARS-CoV-2 infection, including responses associated with inflammation, interferon-beta, and microbial defense mechanisms. The complement and coagulation cascades pathway and representative proteins were activated and upregulated in aged mice after infection. This dual phenomenon of attenuated antiviral response alongside increased activation of the complement and coagulation cascades may contribute to severe COVID-19 in aged mice, suggesting that targeting the complement pathway is a promising therapeutic strategy for managing the elderly population with COVID-19.

## Author contributions

Li Ma: Writing -Original Draft, Writing -Review & Rditing, Data Curation, Formal Analysis, Visualization. Xian Lin: Investigation, Writing - Review & Rditing, Validation. Meng Xu: Data Curation. Xianliang Ke: Investigation. Di Liu: Project Administration. Quanjiao Chen: Writing - review & editing, Funding Acquisition, Project Administration, Resources. All authors have read and agreed to the published version of the manuscript.

## Acknowledgments

We thank Prof. Meilin Jin (Huazhong Agricultural University) for kindly providing the MAV. We also thank Tao Du and Jin Xiong of National Biosafety Laboratory, Wuhan for their essential support. We are grateful for the technical support of the Institutional Center for Shared Technologies and Facilities of Wuhan Institute of Virology, CAS (Center for Instrumental Analysis and Metrology and Center for Experimental Animals). We would like to thank Editage (www.editage.cn) for English language editing.

## Conflict of interest

The authors declare that they have no conflict of interest.

## Data availability

The RNA-seq raw and processed data were deposited in the National Center for Biotechnology Information Gene Expression Omnibus database (series accession no. GSE274784). Proteomics raw and processed data from the LC–MS analysis were uploaded in ProteomeXchange (identifier no. PXD055174) (Please log in the reviewer account first and click on" Review Submission". The original data will be public after the official publication of this article). We have included a pre-processed data table containing a matrix of molecular counts for all samples profiled in our study. The data and code for analysis is shown here: https://github.com/LiMahhhh/mice_bulk_RNA_protein_analysis.

## Funding

This work was supported by the National Key Research and Development Program of China (NKPs) (2022YFC2604101).

## Ehics statement

All animal experiments were conducted in an Animal Biosafety Level 2 (ABSL-2) laboratory and were approved by the Animal Experimental Ethics Committee of the Wuhan Institute of Virology, Chinese Academy of Sciences (approval number: WIVA04202107).

---

**Supplementary Fig. S1.**
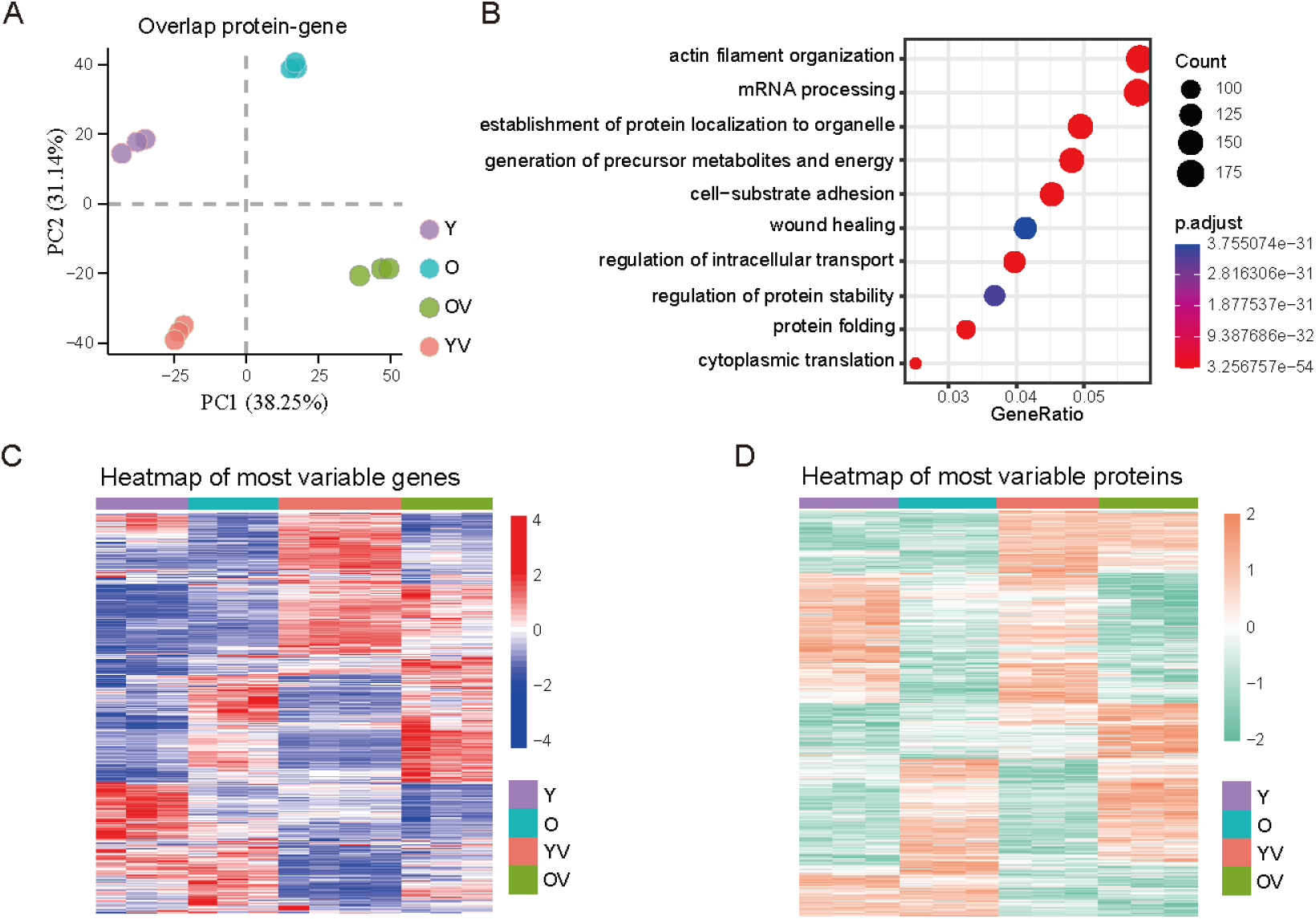
Integrative Analysis of Proteomic and Transcriptomic Data. (A) PCA plot illustrating variance in groups with overlapped proteins/gene expression profiles. (B) Bubble plot displaying GO enrichment of genes from overlapped proteins/genes expression profiles. (C) and (D) Heatmap showing scaled expression of highly variable genes and proteins among groups Y, O, YV, and OV.

**Supplementary Fig. S2.**
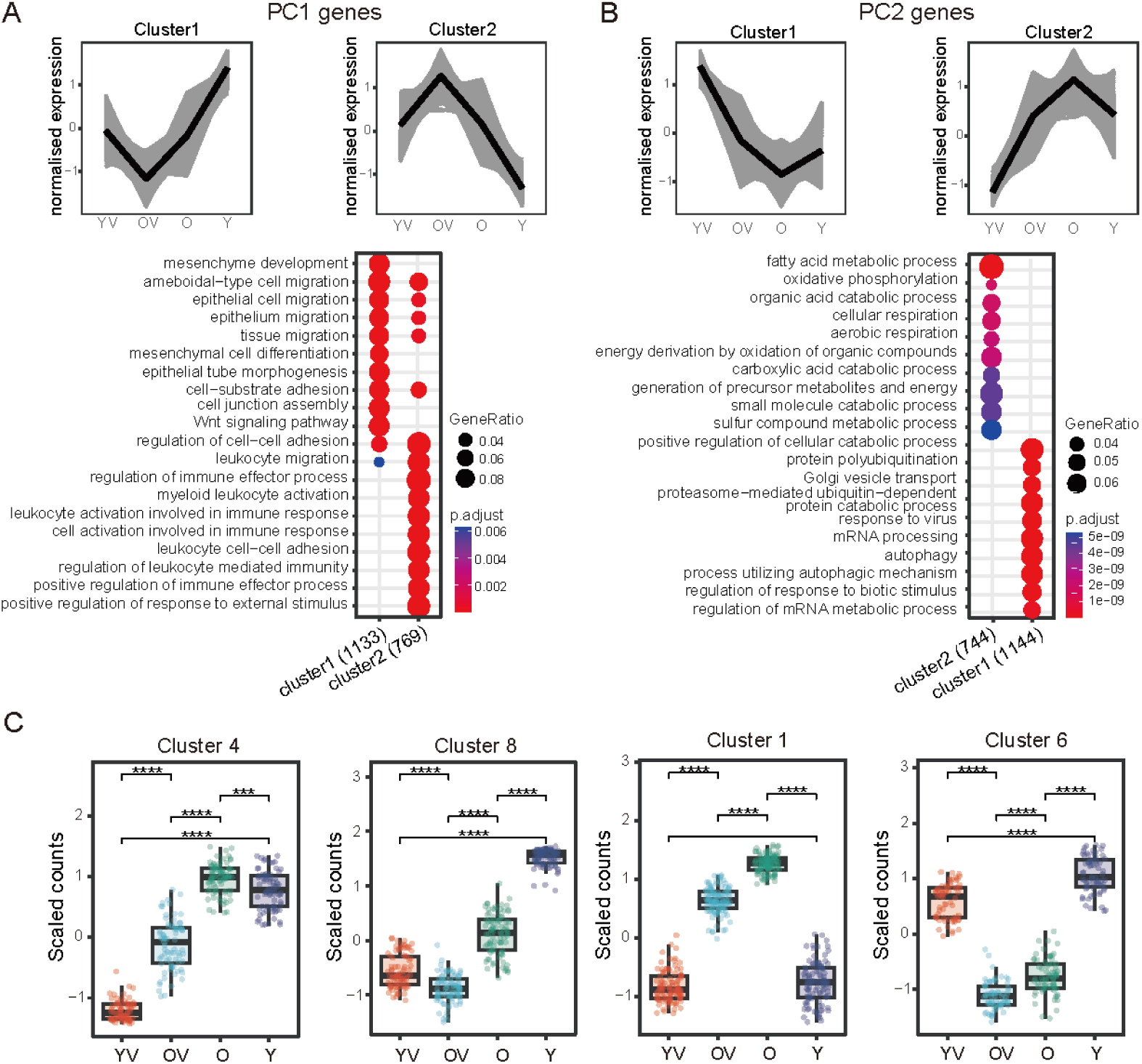
Differential biological processes and clusters associated with age and virus infection. (A) and (B) Expression of the top 50 genes (upper panel) and GO enrichment analysis of the top 1000 genes (lower panel) contribution to Principal component (PC) 1 (A) and PC2 (B). Line graphs depict normalized gene expression (grey) and the average expression levels (black). The bubble chart highlights the key biological processes. (C) Box plots illustrating the scaled expression levels of selected gene clusters (4, 8, 1, and 6) among groups Y, O, YV, and OV. Asterisks indicate statistical significance levels (****, p<0.0001).

**Supplementary Fig. S3.**
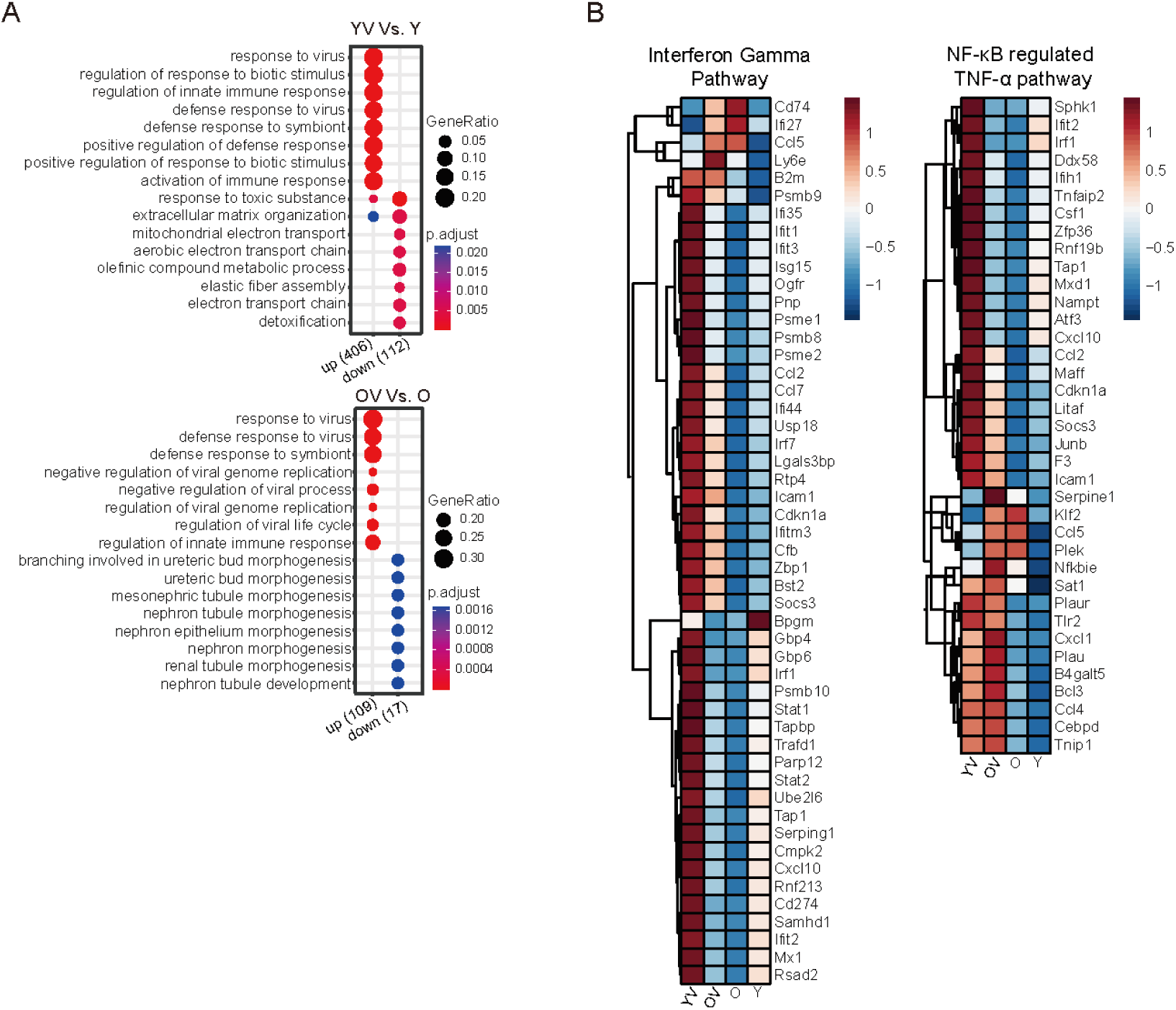
DEG-linked GO network. This network illustrates the association of significantly altered biological function GO terms (pie charts; adjusted p < 0.05) with DEGs (small nodes) across comparisons: O Vs. Y, OV Vs. O, YV Vs. Y, and OV Vs. YV. The pie chart size corresponds to the number of genes involved, and linked terms indicate shared genes. Dash lines denote clusters of closely related functions arranged based on topology. Heatmaps display transcript counts for each group, aligned with clusters and DEGs in Fig. 2C.

**Supplementary Fig. S4.**
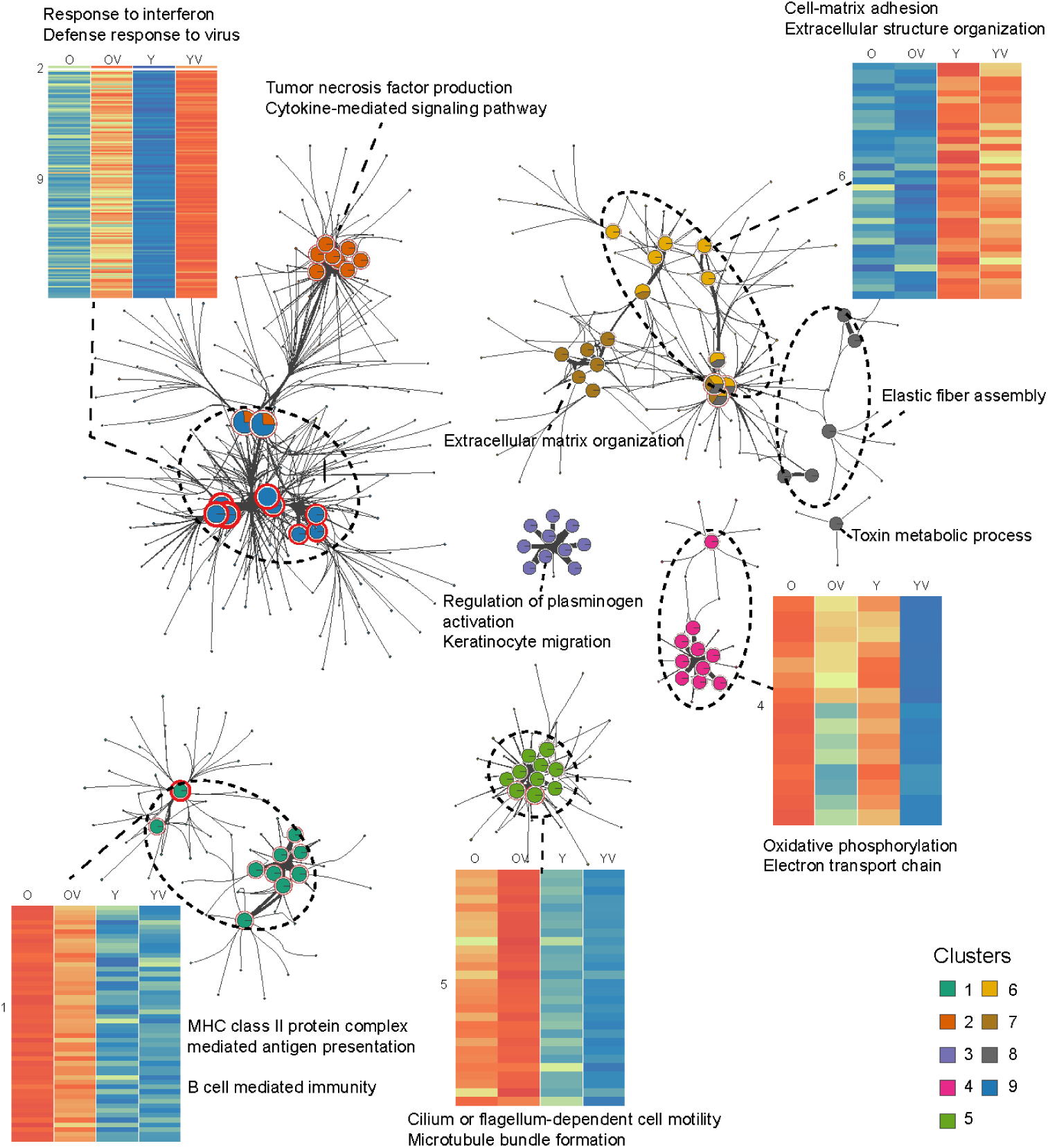
Functional enrichment and pathway activation analysis. (A) Bubble plots illustrate the GO enrichment of the DEGs between YV and Y, as well as OV and O. Bubble size indicates the fraction of DEGs found in the gene set, with color intensity representing statistical significance. (B) Heatmaps showing scaled gene expression of HVGs in the Interferon Gamma Pathway and NF-κB regulated TNF-α pathway among groups Y, O, YV, and OV.

**Supplementary Fig. S5.**
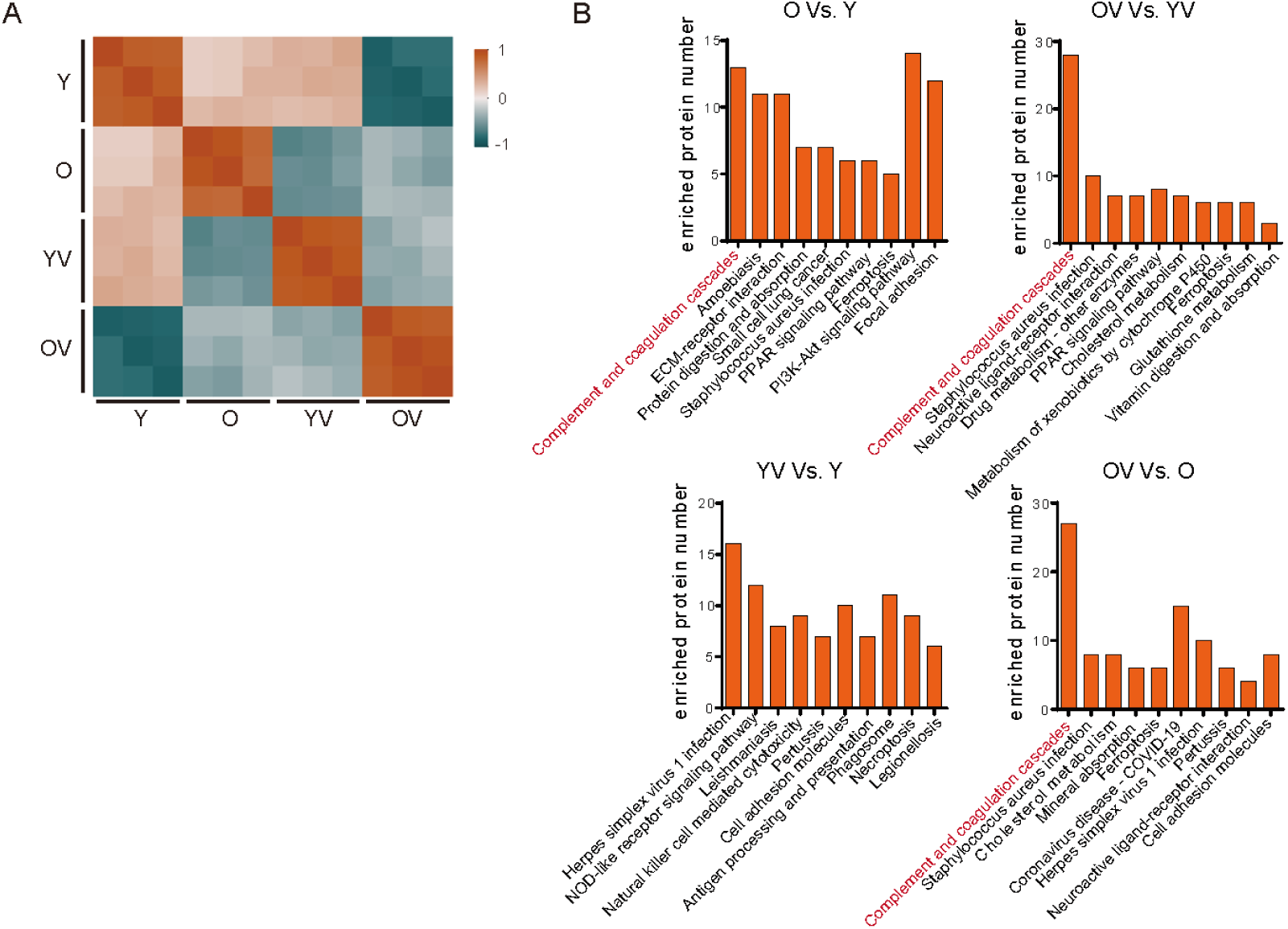
Comparative analysis of proteome. (A) Heatmap illustrating the similarity in protein expression patterns between groups, with color intensity indicating correlation strength. (B) Bar graphs showing protein counts in enriched KEGG pathways across comparisons: O Vs. Y, OV Vs. O, YV Vs. Y, and OV Vs. YV.

## Notes

### Competing Interest Statement

The authors have declared no competing interest.

